# Single-nucleus expression characterization of non-enhancing region of recurrent high-grade glioma

**DOI:** 10.1101/2022.10.10.511639

**Authors:** Kunal S. Patel, Kaleab K. Tessema, Riki Kawaguchi, Alvaro G. Alvarado, Sree Deepthi Muthukrishnan, Akifumi Hagiwara, Vivek Swarup, Linda M. Liau, Anthony Wang, William Yong, Daniel H. Geschwind, Ichiro Nakano, Steven A. Goldman, Richard Everson, Benjamin M. Ellingson, Harley I. Kornblum

## Abstract

**Background:** Non-enhancing (NE) infiltrating tumor cells beyond the contrast-enhancing (CE) bulk of tumor are potential propagators of recurrence after gross total resection of high-grade glioma.

**Methods:** We leveraged single-nucleus RNA-sequencing on 15 specimens from 5 high grade gliomas to compare prospectively identified biopsy specimens acquired from CE and NE regions. Additionally, 24 CE and 22 NE biopsies had immunohistochemical staining for Ki67 to identify proliferative cell burden.

**Results:** Tumor cells in NE regions are enriched in neural progenitor cell-like cellular states, while CE regions are enriched for mesenchymal-like states. These NE glioma cells have similar proportions of proliferative and putative glioma stem cells relative to CE regions, without significant differences in % Ki67 staining. Tumor cells in NE regions exhibit upregulation of genes previously associated with lower grade gliomas. Cell-, gene-, and pathway-level analyses of the tumor microenvironment in the NE region reveal relative downregulation of tumor-mediated neovascularization and presence of cell-mediated immune response, but increased glioma-to-non-pathological cellular interactions.

**Conclusions:** This comprehensive analysis illustrates differing tumor and non-tumor landscapes of CE and NE regions in high-grade gliomas, highlighting the NE region as an area harboring likely initiators of recurrence in a pro-tumor microenvironment and identifying possible targets for future design of NE-specific adjuvant therapy.

**Key Points:** Significant proliferating tumor burden exist in non-enhancing regions of glioma; non-enhancing regions have unique tumor and non-tumor expression properties

**Importance of Study:** Standard of care treatment for glioblastoma relies on visualization of tumor via contrast-enhanced magnetic resonance imaging. However, non-enhancing regions harbor tumor cells that should be targets for adjuvant therapy given these regions are not resected in surgery. To begin addressing these infiltrating non-enhancing tumor cells, we thoroughly characterize the tumor and non-tumor microenvironment of non-enhancing regions in high grade gliomas. Understanding the total tumor burden, proliferating tumor ratio, and presence of putative glioma stem cells may help design adjuvant therapies for these unique population of tumor cells. Understanding the non-tumor immune and vascular microenvironment may help target these areas in regards to drug delivery and immunotherapy. Overall, in a disease marked by significant intratumoral heterogeneity, we focus identifying therapeutic strategies for areas not addressed at surgery.

## Introduction

The intervention that contributes most to overall survival in adult glioma remains surgical removal of tumors ^1,2^. The current surgical standard of care is imaging-based resection of the contrast-enhancing (CE) region as depicted by gadolinium contrast-enhanced T1-weighted magnetic resonance imaging (MRI) ^3^, where increased extent of resection ^4^ and lower post-surgical residual enhancing disease ^5^ have been shown to be correlated with improved survival. However, despite total resection of the CE region, high-grade gliomas invariably recur, and this most commonly occurs at the edge of the resection cavity ^6^.

Treatment options for recurrent malignant glioma include repeat resection with adjuvant molecular therapy ^7–9^ and/or immunotherapy ^10^. Given the modest efficacy in randomized trials, further study into treatment for recurrent glioma is needed. Strategies for molecularly targeted therapy for recurrent glioma have largely been developed by studying bulk RNA or protein expression from cell lines or clinical surgical specimens, which, as described above, generally derive from the CE portion of the tumor. This strategy 1) fails to account for cellular heterogeneity and 2) ignores the true targets of adjuvant therapy for glioma: the residual cells beyond the CE region that are not resected. While more recent studies have utilized single-cell sequencing to study intratumoral heterogeneity ^11–18^, they have largely focused on single samples from CE regions, yielding limited insight about infiltrating cells beyond the CE region and their clinical relevance. Furthermore, in addition to understanding infiltrating tumor cells, it is important to understand the non-tumor microenvironment along with tumor-microenvironment interactions in regards to immune response, vasculature, and normal nervous system function ^10^.

In this study, we prospectively sample CE regions and tissue beyond the enhancing edge (non-enhancing, “NE”) from five high grade gliomas: three grade 4 IDH-wildtype glioblastomas, one grade 4 IDH-mutant astrocytoma, and one grade 3 IDH-mutant oligodendroglioma. We couple intraoperative neuronavigation-guided targeting of biopsy specimens with single-nucleus RNA-sequencing (snRNA-seq) to comprehensively characterize region-specific cellular and molecular features. Our experimental design differs from previous single-cell analyses in glioma in that we leverage prospectively determined positional data to model the spatial landscape of cell populations within CE and NE regions, producing an improved ^19^ yield of single-cell data from NE regions of tumor. We perform a comprehensive single-cell characterization of the NE region to assess tumor cell burden, composition of tumor and non-tumor cells, molecular features of tumor and non-tumor cells, and tumor-microenvironment interactions. These data provide an important description of the biological landscape of infiltrating tumor cells and identify potential targets for improved post-operative adjuvant therapy.

## Methods

### Lead contact and materials availability

Please direct requests for further information and/or resources to Harley Kornblum.

### MRI-guided biopsy targets in human gliomas

Patients with recurrent high-grade glioma were included in this study. We chose recurrent glioma given its clinical burden and limited single-cell analysis of recurrent glioma relative to primary glioma ^13^. Each patient had a single previous resection with adjuvant temozolomide and radiation therapy only. Patients referred to UCLA Neurosurgery for intrinsic supratentorial brain tumors underwent 3 Tesla thin-cut pre-operative MRI (3T Siemens Prisma, Siemens Healthcare, Erlangen, Germany) with and without gadolinium contrast (gadopentetate dimeglumine; Magnevist; Bayer HealthCare Pharmaceuticals) according to the international standardized brain tumor imaging protocol ^20^ within 1 week of surgery. Images of the following sequences were downloaded: T1 three-dimension magnetization-prepared rapid gradient echo (MP-RAGE) with and without contrast. Using AFNI, a software for analysis and visualization of functional magnetic resonance neuroimages ^21^, three 5mm × 5mm × 5mm spherical biopsy targets were selected per patient by a multidisciplinary team (neurosurgery, neuroradiology, and neurooncology) based on feasibility and anatomic landmarks in order to minimize surgical inaccuracy from brain shift. Biopsy targets were then transferred to Brainlab Curve (BrainLAB AG, Munich, Germany), a surgical neuronavigation software, for intraoperative image guidance and tissue acquisition. There was no change to surgical and post-surgical standard of care therapy ^22^ for all patients. Specimens were immediately fresh frozen for nuclear isolation or paraffin-embedded for immunohistochemistry.

### Immunohistochemistry analysis

MRI-targeted biopsy specimens were collected from 17 patients with malignant glioma, yielding 22 specimens from NE regions and 24 from CE regions. Specimens were paraffin-embedded and stained with anti-Ki67 antibody.

Image based quantification was done using QuPath ^23^ to calculate percentage of sample pixels with positive staining. Samples were compared using independent samples t-tests as well as paired samples t-tests in the five patients from whom both CE and NE specimens were acquired.

### Single-nucleus RNA sequencing (snRNA-seq) of glioma specimens

Single nuclei were isolated from frozen tumor specimens using iodixanol-based density gradient centrifugation and submitted to UCLA Technology Center for Genomics and Bioinformatics for library preparation and sequencing (*Supplemental Methods, isolation of single nuclei from tumor specimens)*. Single-nucleus cDNA libraries were generated using the Chromium Single Cell 3’ v3 kit (10x Genomics) and sequenced at 600 million reads/library and 2×50 base pairs using the NovaSeq 6000 S2 platform (Illumina). Raw reads were demultiplexed and Cell Ranger (10x Genomics) was used for alignment (human genome GRCh38), filtering, barcode identification, and counting of unique molecular identifiers, resulting in a feature-barcode (i.e., gene-cell) matrix for each biopsy specimen (*Supplemental methods, Single nucleus library preparation, sequencing, read alignment*)..

### SnRNA-seq bioinformatic processing and analysis

Detailed methods are described in supplemental methods. Pre-processing, integration, and clustering was conducted using the R package Seurat ^24–26^ (*Supplemental Methods, SnRNA-seq integration and clustering)*. To identify malignant cells in each patient dataset, we developed a multi-step cell classification approach integrating both gene expression profiles and iterative prediction of copy number alteration (CNA) profiles (*Supplemental Methods, Streamlined CNA inference tool*). To identify non-malignant cell types, we developed an improved tool for identifying cell types based on canonical cell type markers (*Supplemental Methods, Cell type marker visualization tool*). We used the integrated model of glioblastoma malignant cellular states ^27^ to classify glioma cellular states (*Supplemental Methods, Molecular classification of glioma cellular states*) and cycling cells (*Supplemental Methods, Identification of cycling cells*). We used previously described definitions of putative glioma stem cells ^28–34^ to label malignant cells as such (*Supplemental Methods, Identification of putative glioma stem cells*). Cell-cell communication analysis was performed using the CellChat package^35^ (*Supplemental Methods, Cell-cell communication analysis*).

### Estimating extent of tumor burden

Cell type abundances from single-cell sequencing data were used to calculate the tumor cell burden at different biopsy sites. Tumor cell number was normalized by the non-tumor oligodendrocyte number to control for total cells sequenced, which varied per sample. Segmentations of contrast-enhancing lesion of tumor were calculated using previously described methods ^36^. Using AFNI ^21^, this segmentation was minimized to obtain an ROI at the volumetric center of the lesion. The distance between the prospective biopsy ROI and this center of lesion ROI was calculated. The tumor:oligodendrocyte ratio was plotted against this distance. A quadratic model was implemented ^37^:

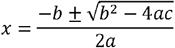

and x solved to identify the Y asymptote as the predicted distance from center when cell ratio approaches 0. A spherical heatmap ROI with radius r equal to this distance was created and overlayed on T1 with contrast images.

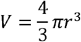

### Statistics

Statistical analysis was carried out in GraphPad Prism (GraphPad Software, San Diego, CA, USA). Independent and paired t-tests and chi-square tests were used to evaluate differences in tumor and non-tumor cell quantities, cellular states, cell cycling proportions and % positive Ki-67. Differential gene expression and pathway enrichment analyses underwent correction for multiple comparisons using the native functionality of the Seurat and fgsea packages, respectively. Significance was defined using an alpha level of 0.05 except as noted in figures.

### Study Approval

This study was approved by the UCLA IRB #10-000655 and #14-001261. All patients provided informed consent for all medical and surgical procedures and involvement in research studies.

### Data availability

Sequencing data will be made available to the public through a GEO submission. Processed data objects and custom scripts for downstream analyses and figure generation will be shared through a GitHub repository at the time of publication (and is available upon request for the editorial process).

## Results

### Analysis of human recurrent gliomas using single-nucleus RNA sequencing

We prospectively identified biopsy targets using pre-operative MRI in 5 patients undergoing surgery for high-grade glioma (Table 1). We selected two biopsy targets from the contrast-enhancing (CE) region of each tumor and one from the non-enhancing (NE) region, located 5-20mm from the CE edge (Fig. 1A). Clinical pathology and fluorescence in situ hybridization identified five high-grade gliomas (three grade 4 IDH-wildtype glioblastomas, one grade 4 IDH-mutant astrocytoma, one grade 3 IDH-mutant and 1p/19q co-deleted oligodendroglioma) (Table 1). We refer to these tumors as Gr4-GBM, Gr4-AST, and Gr3-ODG, respectively ^38^. To analyze these tumors, we performed single-nucleus RNA-sequencing (snRNA-seq) of 15 biopsy specimens (Fig. 1B), yielding 32,914 individual transcriptomes that passed rigorous quality control and were included in subsequent analyses (Table 1, Fig. 1C).

**Table 1.**
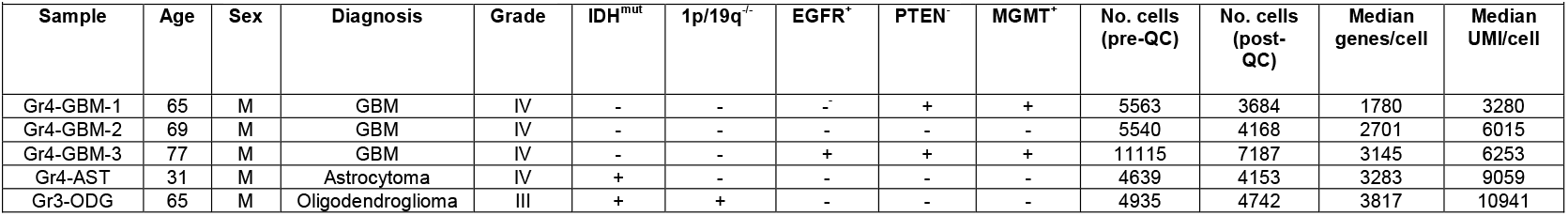
Clincal, Pathological, and Sequencing Characterisics of Included Samples

**Figure 1.**
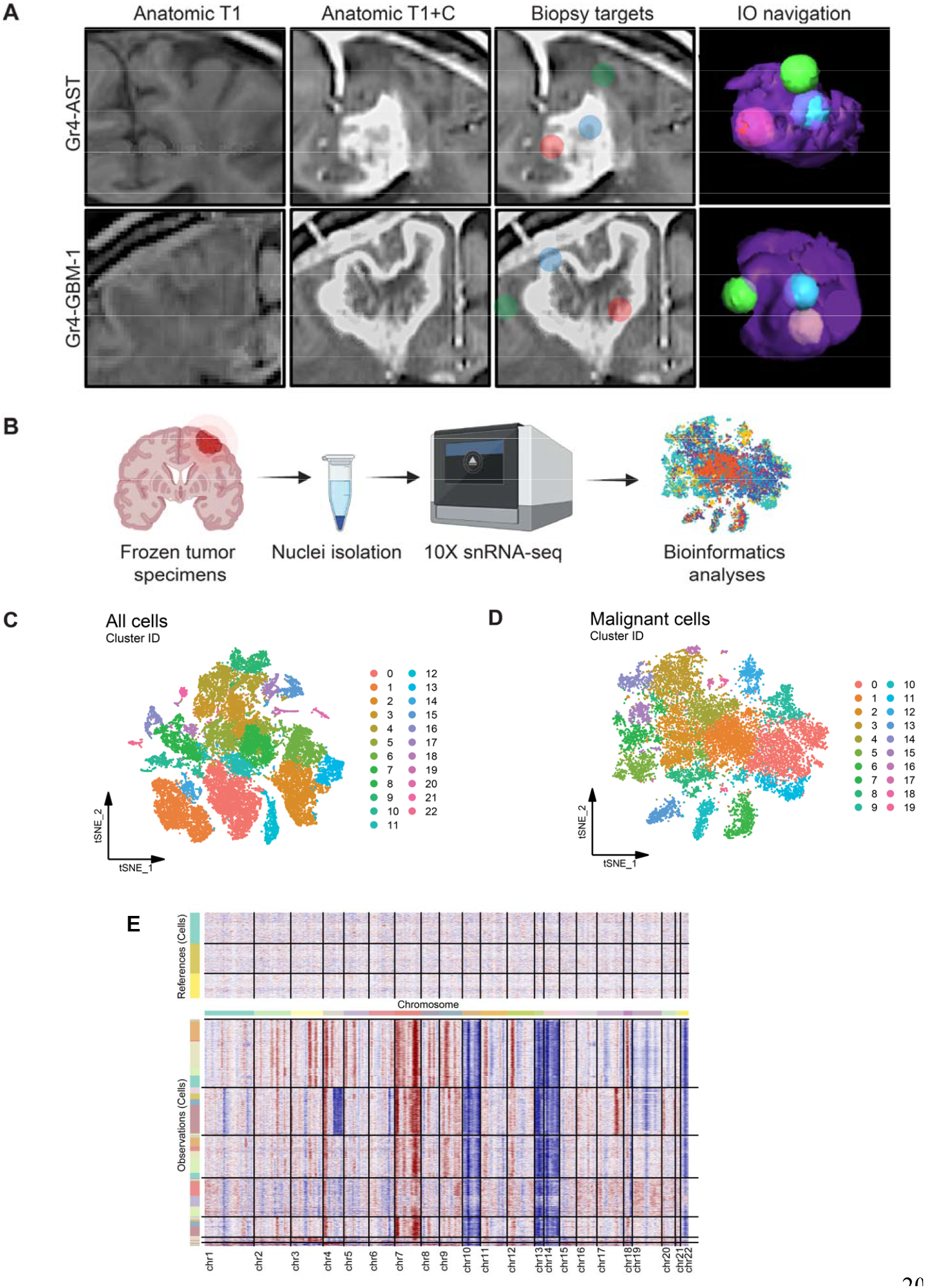
Single-nucleus RNA-sequencing analysis of MRI-guided glioma biopsy targets. **(A)** Preoperative targeted MRI: T1-weighted images without contrast (T1), with contrast (T1+C), prospective biopsy targets, and intraoperative (IO) neuronavigation renderings of contrast-enhancing (blue and red) and non-enhancing (green) samples. **(B)** Single-nucleus RNA-sequencing (snRNA-seq) experimental workflow **(C-D)** 2-dimensional t-distributed stochastic neighbor embedding (t-SNE) plots showing the integrated snRNA-seq dataset with all sequenced nuclei **(C)** and those identified as malignant **(D)**, and example of copy number variation analysis **(E**).

### Significant tumor burden and altered malignant cell composition in the NE regions of high-grade gliomas

We used MRI-based positional information to compare the contrast-enhancing (CE) and T2 hyperintense non-enhancing (NE) regions (Fig. 2A). We first assessed tumor burden in the NE region by quantifying malignant cell proportions, revealing a notable malignant cell burden present in the NE region in all patients (15.3-60.6% of NE cells; NE / CE malignant cell ratio of 0.58-1.13; Fig. 2B). Given that malignant cells comprised a notable proportion of the NE regions, we sought to model the spatial extent of tumor burden on MR imaging. Using tumor cell quantity from three locations in two tumors, we applied a quadratic model (see Methods) to extrapolate the distance from the center of the tumor to where the tumor burden would approach 0 (Fig. 2C). This model predicted tumor cell presence 4.94 cm and 2.31 cm, respectively, from the center of the tumor, and a spheroid with this radius was constructed and overlaid on the corresponding pre-operative MR image (Fig. 2D). The predicted tumor edge was beyond the CE region by 0.34-2.45 cm. While the invasion and growth patterns of glioma are complex, with cases of multifocal and distant lesions ^39^, our data suggest a consistent presence of marked and potentially clinically significant tumor burden immediately surrounding the CE bulk.

**Figure 2.**
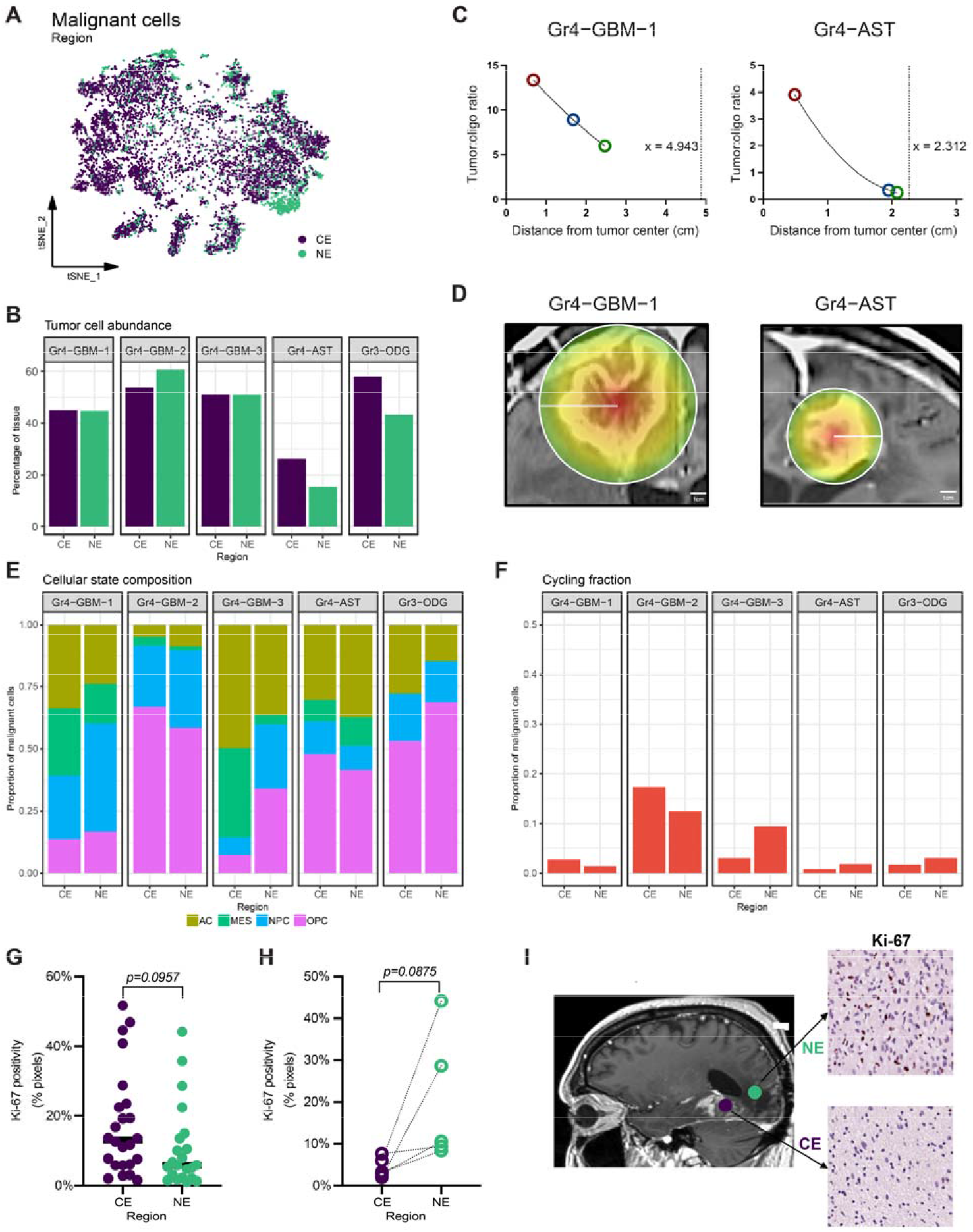
Characterizing tumor burden and composition in the non-enhancing region. **(A)** t-SNE plot of malignant cells from all patients, colored by biopsy region. **(B)** Quantification of tumor burden (proportion of all cells that are malignant) in the CE and NE regions of each tumor. **(C-D)** Tumor burden relative to distance from the tumor center was modeled for Gr4-GBM-1 and Gr4-AST. **(C)** Fitting the tumor cell:oligodendrocyte ratio to a quadratic curve (solid line), with the dotted line representing predicted distance (in centimeters) from the tumor center where tumor burden approaches 0. **(D)** Pre-operative MR scan with superimposed heatmap of predicted tumor burden. **(E-F)** Malignant cellular state composition **(E)** and cycling cell fraction **(F)** in the CE and NE regions of each tumor. **(G-H)** Quantification of immunohistochemistry staining for Ki-67 stratified by region using non-paired **(G)** and paired **(H)** biopsy samples. Statistical comparisons were performed using independent and paired t-tests, respectively. **(I)** Representative example of regions of Ki-67 staining in CE and NE regions.

We then examined malignant cellular states as described previously ^27^ across regions and noted significant heterogeneity both in terms of cellular state composition (Fig. 2E). NE regions harbored a higher NPC-like proportion (mean +14%, paired t-test p=0.03) and trended towards lower MES-like proportion (mean -15.0%, paired t-test p=0.12), while AC- and OPC-like proportions were not consistently different. There was a cycling fraction in the NE regions of all tumors (Fig. 2F), with no significant difference between regions (mean +0.51% in NE, paired t-test p=0.792). To further characterize tumor cell cycling via immunohistochemistry, we obtained image-guided biopsies from an additional cohort of 17 patients with high-grade glioma and analyzed a total of 22 tissue specimens from NE regions and 24 from CE regions (Table 2). Quantification of Ki-67 staining showed a wide distribution in both regions, including several NE samples with high Ki-67 positivity, but no significant difference between regions (p=0.0957, unpaired t-test; Fig. 2G). We also analyzed paired samples from five patients with both regions sampled during the same procedure, with again no significant difference between regions (p=0.0875, paired t-test; Fig. 2H-I). Interestingly, we found that the cycling proportion of malignant cells in CE (but not NE) was predictive of tumor burden in NE (p<0.001, Spearman’s rho=1). Together, these data indicate that there are ample numbers of proliferating tumor cells in the NE region, a potential etiology for recurrence.

**Table 2.**
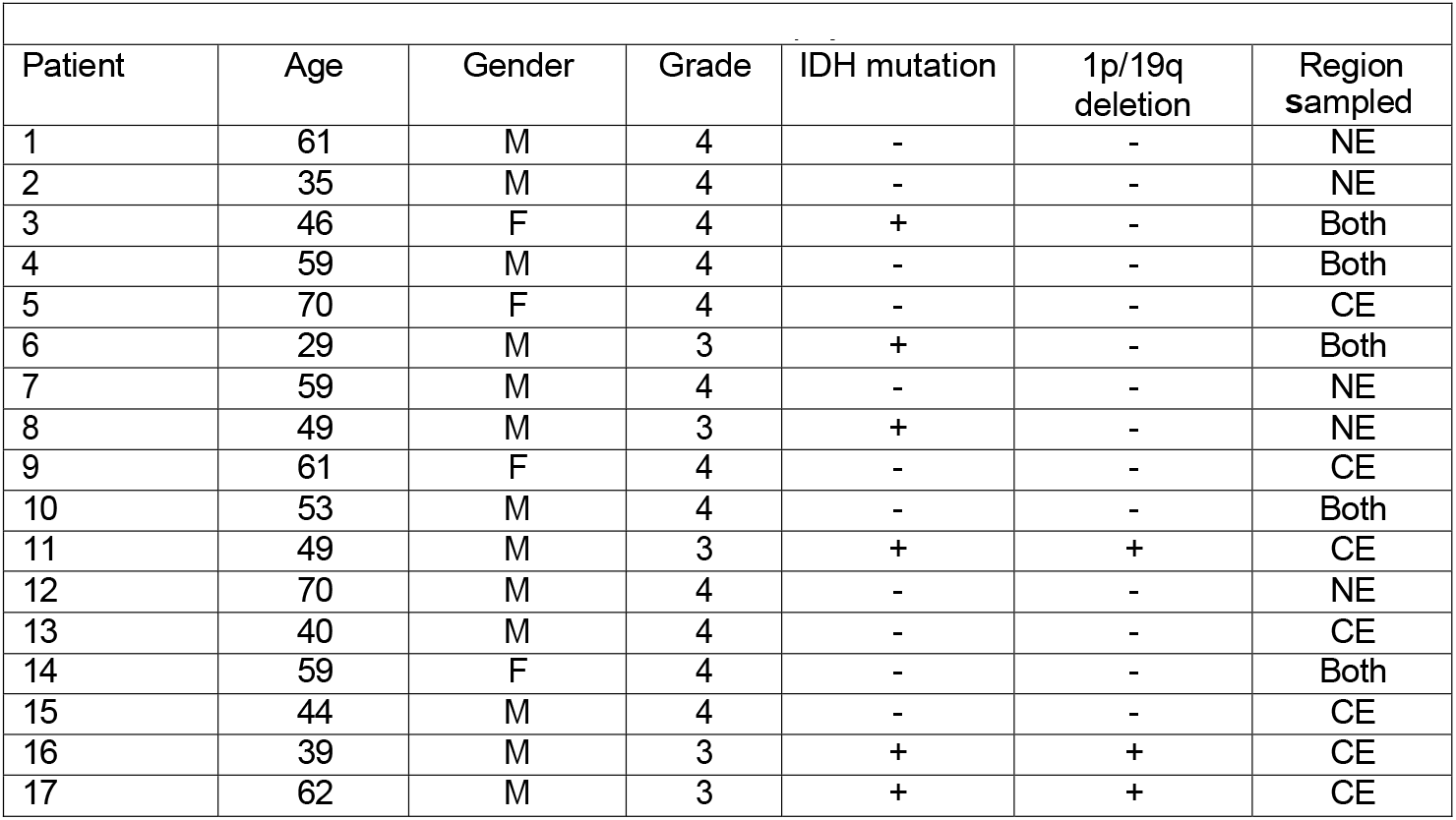
Patient cohort information for localized biopsy immunohistochemistry

### Gene expression and pathway enrichment profiles in malignant cells from NE regions

To investigate regional heterogeneity in malignant cell transcriptomes, we identified differentially expressed genes (DEGs) between regions (Fig. 3A-B). Top NE-enriched DEGs (change > 25% and adjusted p < 1e^-10^) included genes associated with neuronal/synaptic function (e.g., *TNR, NRXN1, ANKS1B, CADM2, IL1RAPL, PAK3, PLCL1, ERC2, SCN3A, KCN3*), genes upregulated in lower grade gliomas (e.g., *TNR, GALANT13, TMEFF2, PAK3, SPOCK1*), and genes associated with immune-cold phenotypes (e.g., *NRXN1*). Top CE-enriched DEGs included genes associated with metabolism (e.g., *ABCA1, NAMPT*), proliferation/invasion (e.g., *CD44, SPP1, IGFBP7, IGFBP5, RYR3, TNC*), and angiogenesis (e.g., *VEGFA, IGFBP7, TNC*). To further determine functional implications of NE-specific transcriptional patterns, DEGs were then used as input for gene set enrichment analysis (GSEA) with a combination of databases: Gene Ontology Biological Pathways, Canonical Pathways, Oncogenic Pathways, and Transcription Factor Targets. Conserved DEGs in malignant cells across tumors pointed to 686 significantly differentially activated pathways in NE vs. CE (Benjamini-Hochberg adjusted p<0.05). The top 20 NE-enriched pathways (normalized enrichment score > 2.0) were all related to neuronal/synaptic function, while the top 20 CE-enriched pathways included interferon-gamma signaling, growth factors, vasculature development, and extracellular matrix regulation. Taken together, our gene-, pathway-, and population-level analyses show significant differences between malignant cells in the NE and CE regions suggesting CE regions are areas of rapid growth, angiogenesis, and inflammation while NE regions harbor glioma cells with potential non-pathological central nervous system (CNS) cell interactions and a landscape that mimics lower grade gliomas.

**Figure 3.**
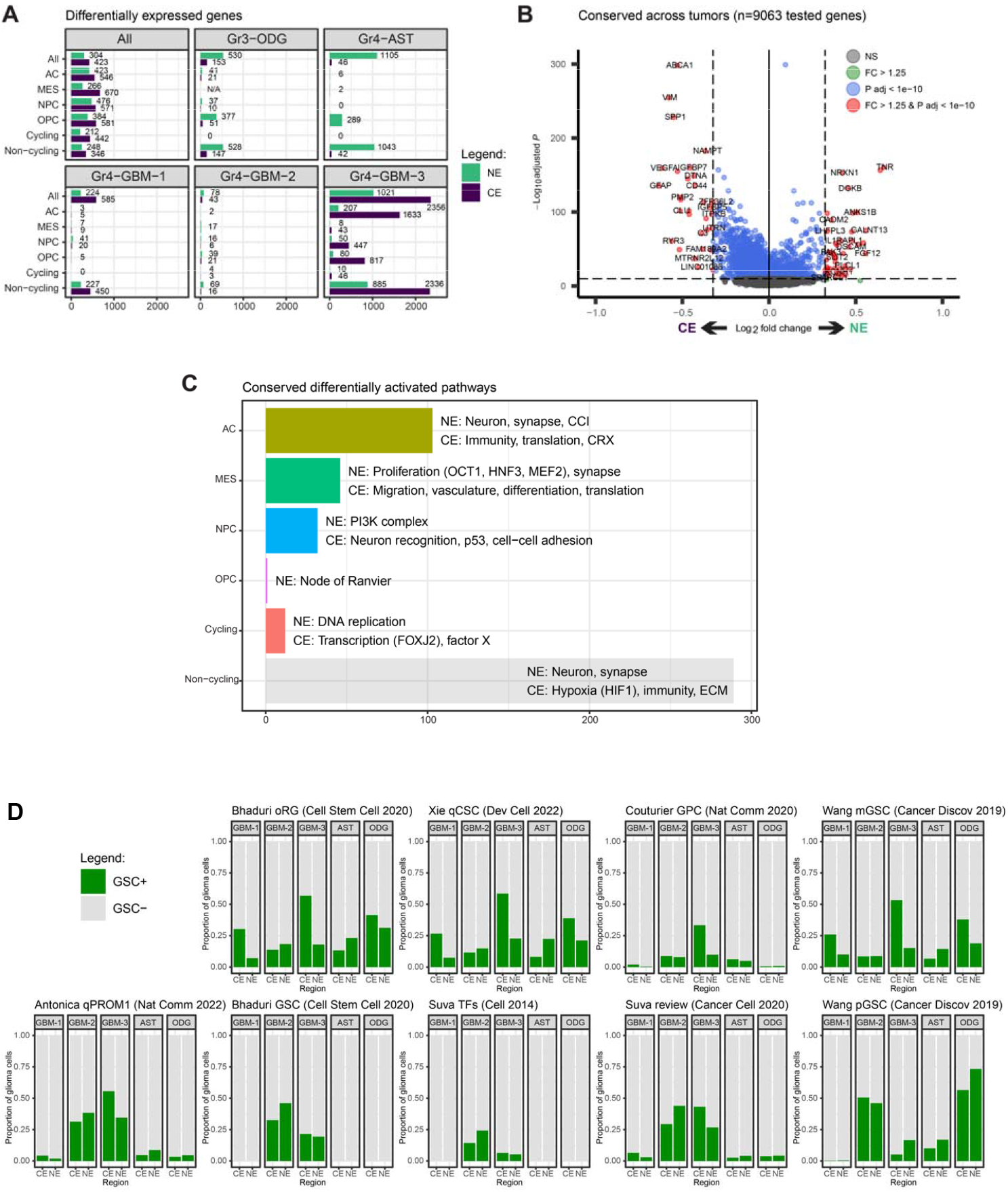
Region-specific molecular features of malignant cells in high-grade glioma. **(A)** Number of NE-enriched and CE-enriched differentially expressed genes (DEGs) in each tumor and malignant cell group. **(B)** Volcano plot showing malignant cell DEGs that are conserved across patients (all cell states combined). **(C)** Number and summary of significantly differentially activated pathways in NE vs. CE identified in each cell group using gene set enrichment analysis (GSEA). DEGs conserved across patients were used as input. The list of tested pathways was a combination of GO Biological Process, Canonical Pathway, Oncogenic Pathway, and Transcription Factor Target databases. **(D)** Relative proportions across regions of putative glioma stem cells (GSCs), identified based on 9 published definitions/gene signatures.

### Regional comparison of putative glioma stem cells

Glioma initiation, progression, invasion, and recurrence may be driven by specific cell populations that are potentially present in residual NE regions. These cells may have certain stem-like properties or specific copy number alterations contributing to their phenotypes ^34^. While putative glioma stem cells (GSCs) have been well studied, there is no universally accepted genomic or transcriptomic definition. To evaluate regional differences in GSC populations, we used nine previously studied “GSC” (for simplicity) gene signatures ^12,16,28,29,32–34,40^ and identified putative GSCs in our dataset (Fig. 4D).There was significant heterogeneity in GSC fraction (mean 0%-38%). Comparing GSC proportion in CE vs. NE yielded no consistent difference (p > 0.05 each). Importantly, all definitions revealed putative GSCs in NE regions. Overall, investigating these glioma populations with previously established roles in tumor initiation, propagation, invasion, and recurrence implicates residual NE regions as harboring critical populations that may serve as targets of adjuvant therapy.

**Figure 4.**
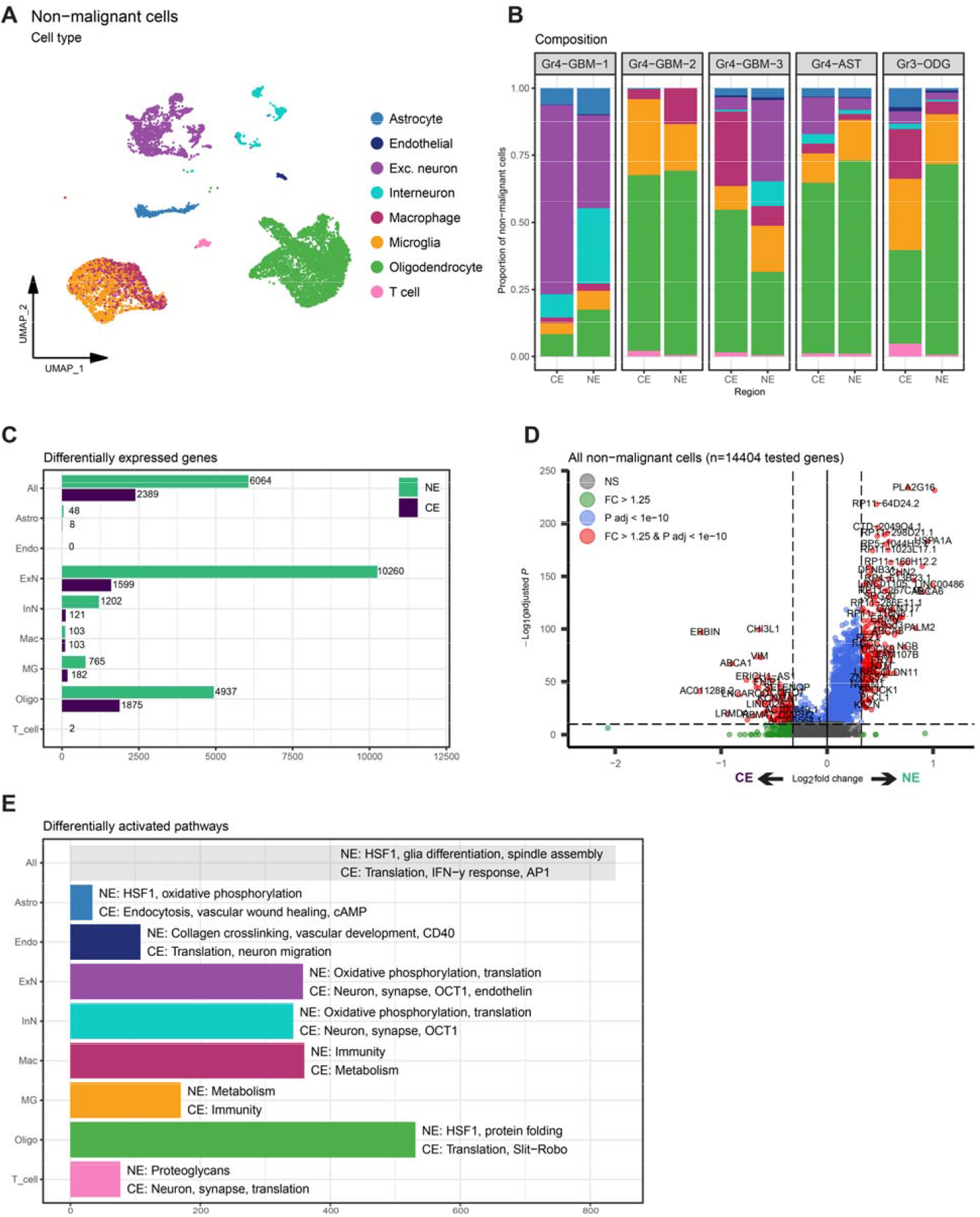
Altered non-malignant cell microenvironment in NE region of high-grade gliomas. **(A)** UMAP representation of non-malignant cells labeled by cell type. **(B)** Cell type composition of non-malignant cells in the NE vs. CE regions of each tumor. **(C)** Number of NE-enriched and CE-enriched differentially expressed genes (DEGs) in non-malignant cell types. **(D)** Volcano plot depicting results of differential expression analysis in all non-malignant cells combined, NE vs. CE. **(E)** Number and summary of cell type-specific GSEA results using the identified DEGs with a combination of the GO Biological Process, Canonical Pathway, Oncogenic Pathway, and Transcription Factor Target databases. *(Astro = astrocytes, Endo = endothelial cells, ExN = excitatory neurons, InN = interneurons, Mac = macrophages, MG = microglia, Oligo = oligodendrocytes, T_cell = T cells)*

### Composition and molecular features of non-malignant cell populations in NE regions

We next focused on regional differences in the tumor microenvironment. Using marker expression patterns and hierarchical clustering, we annotated non-malignant clusters as astrocytes, oligodendrocytes, excitatory neurons, interneurons, endothelial cells, T cells, microglia, and tumor-associated macrophages (Fig. 4A). We first examined cell type composition and noted that oligodendrocytes were the most represented cell type in all tumors except for Gr4-GBM-1, which had a neuronal majority (likely due to the biopsy sites’ proximity to gray matter rather than white matter). This highlights the invasive nature of glioma, with malignant cells integrating into non-pathological CNS tissue. Comparing composition across regions revealed lower T cell proportions in NE for all tumors (mean 0.56% vs 1.94%, paired t-test p=0.14; Fig. 4B). Consistent with previous reports ^19^, the macrophage:microglia ratio was higher in the CE regions of all tumors except Gr4-GBM-2. There were no clear regional patterns in other cell type proportions.

We examined global and cell type-specific regional transcriptomic differences in non-malignant cells by identifying differentially expressed genes (DEGs) in NE vs. CE in the integrated dataset (Fig. 4C-D). In oligodendrocytes, heat shock proteins (e.g., *HSPA1A, HSPH1)* and the metabolic gene *ENOX2* were NE-enriched, while *VEGFA* was CE-enriched. In neurons, genes associated with hypoxic conditions (e.g., *NGB*) and increased metabolism (e.g., *MT-ATP6, MT-CO1, MT-CO2, MT-CYB*) were NE-enriched, while genes related to cell adhesion (e.g., *CADM2, LAMA2, CDH18*) and neuronal function (e.g., *DLG2, NRG3, CSMD1, PARK2*) were CE-enriched. In microglia, heat shock protein genes (e.g., *HSPA1B, HSP90AA1, HSPH1*) were NE-enriched, and a gene associated with monocyte-to-macrophage differentiation (*CPM*) was CE-enriched. In macrophages, genes associated with interaction and reorganization of the extracellular matrix (e.g., *TNS1, VIM, TRIO*) were CE-enriched. Lastly, genes involved in blood-brain barrier (e.g., *CLDN11, PDGFRb*) were NE-enriched in non-tumor cells.

We also assessed regional enrichment of functional pathways in non-malignant cells using GSEA, (Fig. 4E). In oligodendrocytes, we observed upregulated HSF1 activity and downregulated Slit/Robo signaling ^41^ in NE. In neurons, we observed upregulation of oxidative phosphorylation and translation and downregulation of endothelin and neuron/synapse terms in NE. Evaluation of microglia demonstrated upregulation in metabolism and downregulation in immune function in NE, with macrophages interestingly exhibiting the opposite pattern. These cell-, gene-, and pathway-level analyses identify unique features of the non-tumor microenvironment in NE regions, highlighted by decreased T cell fractions, possible region-specific myeloid phenotypes, and markers of cellular stress (e.g., HSF1) in multiple cell types. Furthermore, these data suggest altered regulation of vascular integrity in NE regions, which may explain the limitation of gadolinium leakage from affected blood vessels and therefore lack of contrast enhancement in this region ^42^.

### Cell-cell communication networks in the NE region exhibit differential functional wiring and information flow

Crosstalk between malignant cells and normal cells is a hallmark feature of cancer and is critical to tumor growth, invasion, and recurrence. To analyze cell-cell interaction (CCI) networks, we utilized CellChat ^43^, to build a CCI atlas for each tumor to examine general principles of cellular communication. The five high-grade gliomas were analyzed individually, with malignant and non-malignant cells combined. We identified between 112-150 putatively active CCI pathways in each tumor out of 223 total pathways in the database. A total of 160 unique pathways were identified; 96 were detected in all tumors. We performed differential analysis to identify shared and patient-specific CCI differences between NE and CE (Fig. 5A). Communication strength (i.e., probability) was higher in NE for four of the five tumors (Fig. 5B). In all tumors, glioma to oligodendrocyte interactions were the upregulated in NE relative to CE (Fig. 5C). In regards to specific communication pathways, CD40 signaling was CE-enriched in all tumors, NT and LIGHT/TNFSF14 were CE-enriched in four tumors (Fig. 5D), while PVR/CD155 (invasion/infiltration) and MAG (myelin interaction) were NE-enriched in four tumors (Fig. 5E). Additional conserved CE-enriched pathways included immune (IL1, IL10, IL4, TIGIT) and endothelial (CDH5) mediators, and conserved NE-enriched pathways included heparin binding growth factor pathways (MK, PTN) and microglial attraction (GDNF). Together, these findings highlight global and cell type-specific differences in the cellular communication landscape of NE regions.

**Figure 5.**
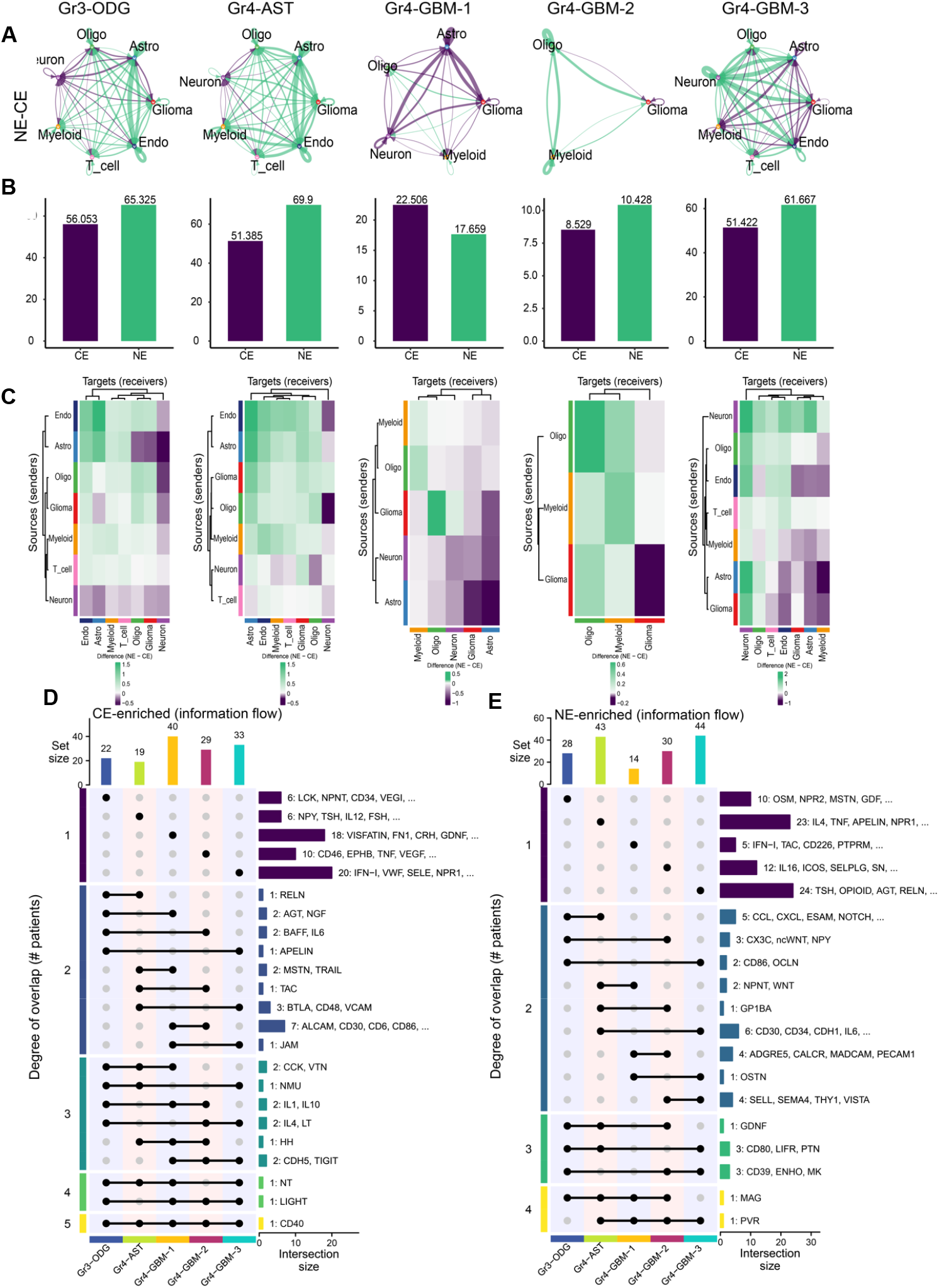
Regional differences in cell-cell interaction strength at the cell type level. **(A)** Differential interaction strength in NE vs. CE for individual cell-cell interactions. Colors correspond to the region with stronger signaling, and line segment thickness corresponds to magnitude of regional difference. **(B)** Total interaction strength between NE and CE networks. **(C)** Heatmap of differential interaction strengths in NE vs. CE among all possible cell type pairs, with rows being senders and columns being receivers of communication. Row and column dendrograms indicate similarity based on hierarchical clustering. UpSet plots depicting overlap of differential information flow enriched in **(D)** CE and **(E)** NE regions. Dots in each row indicate which tumor (degree of 1) or tumors (degree of 2+) comprise each group, and bars on the right indicate how many and which pathways are present exclusively in each group (i.e., pathways can only be present in one row). Bars on the top indicate the number of regionally heterogeneous pathways in each tumor.

## Discussion

In this study, we leveraged single-nucleus RNA-sequencing and MRI-guided biopsy target selection to dissect spatial phenotypes in high grade glioma. While prior studies have characterized cells in both NE and CE regions ^19,44^, our approach has the advantage of being able to capture sufficient numbers of cells from each region to ascertain regional differences. With this in mind, we focused on characterizing these infiltrating NE regions on a single-cell level by taking MRI-guided samples of these areas during surgery. We confirmed there was significant tumor burden beyond the CE region in all recurrent high-grade gliomas. Our model predicted that cells may exist multiple centimeters beyond the CE edge, potentially occurring in a portion of FLAIR hyperintensity surrounding the tumor. NE regions had similar levels of cycling cells, and putative GSCs (using several definitions), CE regions, suggesting these residual areas likely harbor propagators of progression and recurrence after gross total resection of CE tumor.

We use cell-, gene-, pathway-, and interactome-level analyses to characterize both infiltrating tumor and microenvironment in the NE region. We found regional differences in cellular state composition that are consistent with previous findings in primary glioma: Minata et al. ^45^ identified higher MES-like cells in CE regions and higher NPC-like cells in NE regions. Differential expression analysis showed upregulation of genes associated with lower grade tumors and downregulation of genes associated with proliferation/invasion in NE regions. Furthermore, the cycling cells of NE regions were almost exclusively OPC-like and NPC-like. These data suggest potential relationships between cellular states and MRI findings in that NPC-/OPC-like cells correlate with non-enhancing characteristics and lower grade biology, while MES-/AC-like cells correlate with contrast-enhancing characteristics and higher-grade biology. In regards to the tumor microenvironment, we highlight 1) low levels of immune cells and relatively decreased tumor necrosis factor related pathways (e.g., CD40, LIGHT, TIGIT) suggesting a relative lack of immune response in NE regions 2) low levels of endothelial cells with upregulation of BBB proteins supporting lack of contrast extravasation as a biological etiology for NE regions with associated signs of hypoxic stress that may be signaling future neovascularization, and 3) signaling between glioma and non-pathological CNS cells supporting active glioma interaction/infiltration along the NE edge of tumor.

Our study has several limitations. Due to the limited number of tumors in each pathological designation, we cannot confidently draw statistical conclusions between types of tumors. Furthermore, the specimens from the NE region in each tumor were limited to a single site given the need to limit sampling of potential non-pathologic neural tissue, leaving open the possibility that different portions of the NE regions behave differently. Lastly, while studying recurrent gliomas allowed comprehensive characterization and comparison with previous findings in primary gliomas, certain conclusions are limited in their generalizability to primary tumors.

Taken together, this study presents a thorough quantification and molecular characterization of both malignant and non-malignant cell populations and their putative functional roles in the NE region of recurrent gliomas. These findings have significant clinical implications. First, the presence of significant tumor burden with actively cycling cells multiple centimeters from the CE edge suggests that resection of the NE region--when not involving eloquent areas, given the potential maintenance of neuronal function in infiltrative areas--may improve outcomes. These regions are often not addressed by surgery (with standard of care being maximal safe resection of CE regions) or adjuvant radiotherapy (with standard of care being radiation to resection cavity with a 2cm margin). Second, considering the inability to resect entire NE regions in many cases due to functional eloquence, these residual NE cells are likely the culprits of recurrence and therefore should be targeted by adjuvant therapy. The presence of cell-, gene-, pathway-, and interactome-level differences in the NE region compared to CE suggests that improvements in adjuvant therapy should be driven by investigation of NE tissue rather than the bulk of the tumor, as is current standard in glioma studies.

