## Supplemental Methods for "Single-nucleus expression characterization of non-enhancing region of recurrent high-grade glioma"

*Isolation of single nuclei from tumor specimens*

Single nuclei were isolated from all frozen tumor specimens using a protocol based on Krishnaswami et al. (1) that employs iodixanol-based density gradient centrifugation. Specimens were cut into small pieces, homogenized on ice using a Dounce tissue grinder, filtered using a 40uM Corning cell strainer, centrifuged at 1000g for 8 minutes at 4C in Eppendorf tubes, subjected to an iodixanol (OptiPrep; STEMCELL Technologies) gradient of 50% vs 29%, centrifuged at 13500g for 20 minutes at 4C, resuspended in PBS with 1% BSA, and labeled with the DNA stain Hoechst 33342 (Sigma). Nuclei were then purified based on size and Hoechst fluorescence using a microfluidic chip-based gentle cell sorter (On-Chip Sort; On-Chip Biotechnologies). The recovered nuclei were counted using an automated cell counter (Countess; ThermoFisher) and immediately submitted to the UCLA Technology Center for Genomics and Bioinformatics (TCGB) for library preparation and sequencing.

*Single nucleus library preparation, sequencing, read alignment*

Single nuclei from each tumor specimen were counted again and visualized at the TCGB core facility to evaluate quality. Single nucleus cDNA libraries were then generated using the Chromium Single Cell 3’ v3 kit (10x Genomics) following the manufacturer’s protocol, which consists of loading nuclei into the channels of a 10x Chromium chip, partitioning the nuclei into Gel Beads in Emulsion (GEMs), reverse transcription of RNA within the droplets, breaking the GEMs, amplification, fragmentation, and addition of adaptor and sample index. For single nucleus RNA-sequencing (snRNA-seq), each library was split across multiple lanes of the NovaSeq 6000 S2 platform (Illumina) and sequenced to a depth of approximately 600 million reads per library, with a read length of 2x50 base pairs. The resulting raw reads were then demultiplexed and processed using Cell Ranger (10x Genomics) for alignment to the human genome (GRCh38), filtering, barcode (i.e., cell) identification, and counting of unique molecular identifiers (i.e., transcripts). The output feature-barcode (i.e., gene-cell) matrices contained the quantitative transcriptomic information for all single nuclei isolated from each tumor specimen.

*SnRNA-seq pre-processing, quality control, and analysis*

Detailed methods are described in supplemental methods. Feature-barcode matrices were further processed and analyzed individually using the R package Seurat (2–4) except as noted below. Plots were generated with functions native to Seurat or from the visualization package scCustomize (5). For quality control, genes expressed in fewer than 5 nuclei and nuclei expressing fewer than 500 unique genes were both removed. Specimens with a markedly low average number of unique genes expressed per nuclei were removed from the study. Nuclei with mitochondrial transcript content of 5% or greater or ribosomal transcript content of 2% or greater were removed. Counts were normalized and scaled using SCTransform (6). For each patient, the datasets from all specimens were then integrated using Seurat’s canonical correlation analysis to account for technical variation (3). Dimensional reduction was performed on each patient dataset via principal component analysis (PCA), uniform manifold approximation and projection (UMAP) embedding, and t-distributed stochastic neighbor embedding (t-SNE). Clustering was performed in each patient dataset using Seurat’s default graph-based clustering, and low dimensional plots were generated using the DimPlot function. Malignant cells and subclones were identified using a custom tool that combines marker expression patterns, hierarchical relationships across unbiased clusters, and an iterative sequence of single-cell copy number alteration (CNA) inference analyses to address problematic cells (i.e., incorrect cluster assignment, noisy or unclear CNA profiles) (see Supplemental Methods). Malignant cells were grouped by cellular state (AC-, MES-, NPC-, or OPC-like) and by cycling status using single-cell gene module scores, as reported by Neftel et al. (7) and their respective pathway enrichment profiles were identified with enrichR (8–10). Non-malignant cell types were identified based on marker expression profiles (using a modified tool for improved visualization of marker expression across clusters) and hierarchical clustering, yielding astrocytes, oligodendrocytes, excitatory neurons, interneurons, endothelial cells, T cells, tumor-associated macrophages, and microglia. The integrated dataset for each patient was then split into malignant and non-malignant datasets, which were processed again as above then integrated across patients. The two resulting datasets (malignant cells from all patients, non-malignant cells from all patients) were then prepared for further analysis by dimensional reduction and Seurat clustering as above. Differentially expressed genes (DEGs) were identified using the Seurat functions FindMarkers (within-group) and FindConservedMarkers (shared) and visualized using EnhancedVolcano. Genes expressed in at least 5% of cells were included; significance was defined as a 10% change in expression with p<0.05 (adjusted p-value for FindMarkers, combined p-value for FindConservedMarkers). Gene set enrichment analysis (GSEA) was performed on DEGs ranked by fold change using the fgsea package (11) with a combination of Gene Ontology Biological Pathways, Canonical Pathways, Oncogenic Pathways, and Transcription Factor Targets (12).

*SnRNA-seq integration and clustering*

Pre-processing and quality control were performed as described in the main text. For each patient, the datasets from all biopsy specimens were then combined using Seurat integration(13), which performs canonical correlation analysis (CCA) to align specimens based on matched biological states (i.e., to correct for technical variation across samples). Integration was performed as recommended by the Seurat developers, specifically by selecting integration features, then identifying anchors, then integrating the datasets. The SelectIntegrationFeatures (using 5,000 genes), PrepSCTIntegration, FindIntegrationAnchors (using the first 50 principal components), and IntegrateData functions were used for integration. Each integrated dataset (one per patient) was then prepared for further analysis by dimensional reduction via principal component analysis (PCA), uniform manifold approximation and projection (UMAP) embedding, and t-distributed stochastic neighbor embedding (t-SNE). These were performed using the functions RunPCA, RunUMAP (using the first 30 PCs and embedding into 3 spatial dimensions), and RunTSNE (using the first 30 PCs) functions, respectively. Finally, each patient dataset was clustered based on gene expression profiles with Seurat’s default graph-based clustering that implements the Louvain algorithm for modularity optimization. Clustering was performed via the functions FindNeighbors and FindClusters. Low dimensional graphs were generated using Seurat’s DimPlot function with default parameters except as noted in the figure captions.

*Identification of malignant cells and subclones*

To identify malignant cells in each patient dataset, we developed a multi-step cell classification approach integrating both gene expression profiles and iterative prediction of copy number alteration (CNA) profiles. Our strategy builds on previous approaches and aims to improve accuracy and reduce noise. This pipeline was performed separately for the integrated snRNA-seq dataset derived from each patient.

First, we examined canonical brain cell type marker expression patterns and hierarchical relationships (see *Cell type marker visualization tool* below) across the clusters identified in each patient to assess whether putative normal brain cell types can be readily identified. Clusters whose marker expression profiles did not clearly align with a normal cell type or aligned with multiple cell types were considered potential malignant cell clusters.

Next, we performed an iterative sequence of single-cell CNA inference analyses for each patient dataset (see *Streamlined CNA inference tool* below). CNA inference was first performed without a reference group of cells (i.e., values were compared to the average across all cells) with cells grouped by their original RNA (Seurat) clusters to examine cluster-specific CNA profiles. To resolve the observed CNA heterogeneity within clusters, cells were then grouped by CNA profile via hierarchical clustering. The resulting heatmap groups cells based on their CNA profiles but also displays the original RNA cluster identities, allowing assessment of potential cell type identities using both pieces of information. Canonical glioma CNA events were clearly visualized in each patient, and cells separated based on these events. Moreover, the resulting CNA clusters were mostly composed of cells belonging to similar RNA clusters, so cells were classified as non-malignant if they belonged to putative non-malignant RNA clusters and non-malignant CNA clusters. For each patient, the heatmap was examined to label CNA-based clusters corresponding to myeloid cells (based on amplification in a portion of chromosome 6 containing the human leukocyte antigen (HLA) system), putative malignant cells (based on canonical glioma-specific CNAs), and putative normal cell types (based on both lack of glioma CNAs and correlation with RNA clusters that were previously labeled as likely normal cell types based on marker expression). Groups of cells that could not confidently be labeled were preliminarily designated as unclassified. Cells that appeared to cluster alone or in very small groups on the dendrogram were designated as noise. Prior to the next iteration, myeloid cells (positive for myeloid markers and chr6 amplification) were removed to focus on brain lineages. CNA inference was then repeated for each patient dataset using a normal cell type as reference, both to remove bias from composition differences across tumors and to identify CNA events relative to normal cells rather than to the average cell. Normal oligodendrocytes were selected as the reference cells since they were the only non-malignant, brain derived (i.e., non-immune) cell type present in all patient datasets. Cells were again grouped by CNA profiles and labeled as in the previous step. For each cell, the labels from each of these steps were used to generate a final set of labels with the following potential values: malignant, non-malignant, noise, and unclassified. This iterative approach allowed us to address multiple types of problematic cells: 1) malignant cells that, due to their atypical transcriptomes, belonged to putative normal RNA clusters, 2) normal cells that belonged to putative malignant or unclear RNA clusters, and 3) cells with noisy or unclear CNA profiles, which we detected at each step. The cells labeled as noise or unclassified were removed, and CNA inference was performed again on the final set of malignant cells using oligodendrocytes as reference and with subcluster identification turned on to identify CNA-based subclones (using the Leiden algorithm) (Fig. S1C). This final set of malignant cells was used for the downstream analyses related to the malignant cell compartment. (Note: To confirm non-malignant cell identities, we also compared CNA profiles of non-malignant cell types across tumors, which showed conserved features in each cell type, e.g., chr6 amplification in myeloid cells and chr11 amplification in astrocytes.)

*Streamlined CNA inference tool*

To perform inference of CNA profiles in single cells, we developed a custom wrapper function that streamlines a workflow leveraging the R package inferCNV (14). The purpose of our wrapper function is to make the CNA inference workflow more efficient and user-friendly by automating the laborious set-up procedure required by inferCNV and adding customization options that do not necessitate manual manipulation of the input files. This function first takes a Seurat object (single-cell expression dataset) and automatically generates all of the input files necessary for inferCNV: raw count data for each cell are obtained via GetAssayData (using the “RNA” assay and the “counts” slot); an annotation file with the Seurat cluster names corresponding to each cell is obtained by extracting the cell names and cluster information from the Seurat metadata; and a gene ordering file with the chromosomal location for all genes present in the dataset is generated using the R package biomaRt (15, 16) to access the GRCh38.p13 human genome assembly via Ensembl (17). The function then passes these input files as arguments to the CreateInfercnvObject function to generate an inferCNV object for the dataset being processed, with an additional option allowing the user to exclude certain cell groups (i.e., Seurat clusters) from the inference analysis. Finally, the function passes the inferCNV object to the run function, which performs the rest of the CNA inference analysis that attempts to identify large-scale chromosomal CNAs by assessing smoothed average gene expression across a moving window of genomic positions, followed by prediction of CNA regions and identification of cell clusters based on CNA profiles. For each patient in the study, an inferCNV object was generated with the X, Y, and mitochondrial chromosomes excluded, then analyzed with default parameters except a cutoff value of 0.1, Hidden Markov Model (HMM) prediction with the i3 model, denoising with a filter of 0.12, and 6 threads for parallelization. This yields several outputs, most importantly a heatmap displaying relative expression intensity in each cell (rows) across chromosomal positions (columns) and text files including the HMM CNA predictions across genes/chromosomes for all cells and cell clusters.

*Cell type marker visualization tool*

To assess expression of canonical cell type markers, we developed an improved tool for generating dot plots (e.g., Fig. S5A). This tool is based on Seurat’s DotPlot function and introduces two improved features. First, the user has the option to automatically color code the cell type labels and corresponding marker genes, which is currently not possible using Seurat’s DotPlot function. Second, the cell clusters are ordered by similarity via hierarchical clustering, with the resulting dendrogram shown to the left of the plot. Importantly, this hierarchical clustering procedure considers all detected genes (or any desired subset of genes) rather than only the genes shown in the dot plot, which is what currently available tools use. The output of our tool is a figure containing both a dendrogram and a dot plot, with color coded cell types and expression information (average expression and percentage of cells expressing) for each marker. The two improvements allow for user-friendly visual customization and more relevant representation of the transcriptomic similarities among clusters (via the dendrogram). These plots were used to examine potential cell type identities based on marker expression profiles and the branching structure seen in the dendrogram.

*Subsetting and processing of malignant and normal cells*

For each patient, the integrated expression object containing all cells was then split into two objects, one including malignant cells and the other including non-malignant cells. Pre-processing was repeated for the individual malignant and non-malignant datasets for each patient. Each dataset was slimmed down to the raw, unprocessed counts using the DietSeurat function to retain only the “RNA” assay. Datasets were then processed using SCTransform^47^. The malignant and normal cell objects from all patients were then combined into two integrated objects using Seurat integration (see *SnRNA-seq integration and clustering* above), with one integrated object containing malignant cells from all patients and one integrated object containing normal cells from all patients. (Note: pre-processing and integration was initially performed to combine the specimens from each patient separately, while this section is describing the same workflow applied to combine the datasets across patients.) Each of these two integrated datasets was then prepared for further analysis by dimensional reduction and Seurat clustering as described in *SnRNA-seq integration and clustering* above.

*Molecular classification of glioma cellular states*

Malignant cells were grouped into subpopulations based on the integrated model of glioblastoma malignant cellular states presented in Neftel et al (18). The same classification and visualization workflows were used in this study. Single cells were classified as astrocyte-like (AC-like), mesenchymal-like (MES-like), neural progenitor cell-like (NPC-like), and oligodendrocyte precursor cell-like (OPC-like) based on enrichment scores corresponding to gene lists related to each of these four cell states. The gene lists corresponding to each cell state can be found in Table S2 of Neftel et al. (18). Enrichment scores were generated for each malignant cell using the Seurat function AddModuleScore with default parameters except assay=”SCT”, search=T, and nbin=30. This function computes enrichment of a gene list (i.e., module score) in each cell by calculating the average expression of the genes contained in the gene list and subtracting the average expression of a control gene list. The control gene list is compiled by first binning all analyzed genes into a certain number of bins (in our case 30) based on aggregate expression levels, then randomly selecting 100 bin-mates for each gene in the query gene list, which results in a control gene set that is 100-fold larger than the query list and has a similar distribution of expression levels. Module scores were generated for all malignant cells using the meta-modules defining the AC-like, MES-like, NPC-like, and OPC-like malignant cellular states, and each cell was assigned to the state with the highest score. Two-dimensional graphs based on these module scores (Fig. 2A) were generated by placed each cellular state into one quadrant of the plot by computing the strength of score separation based on two groupings: OPC-like/NPC-like vs AC-like/MES-like (y-axis) and AC-like/OPC-like vs MES-like/NPC-like (x-axis). The equations used to compute the magnitudes of the x and y coordinates for each cell were as follows:

NPC-like/OPC-like cells:	x = log_2_(|NPCscore – OPCscore| + 1)

AC-like/MES-like cells:	x = log_2_(|MESscore – ACscore| + 1)

All cells:	y = maximum(OPCscore, NPCscore) – maximum(ACscore, MESscore)

The x values were then multiplied by -1 for OPC-like and AC-like cells. This placed the OPC-like cells in the top left quadrant, NPC-like in the top right, AC-like in the bottom left, and MES-like in the bottom right. Cells were colored according to their cellular state assignment or other grouping variables as noted in the figure legends. The top enriched pathways in each cellular state and in cycling cells (Fig. 2b; see *Identification of cycling cells* below) were identified using enrichR (8-10) via the DEenrichRPlot function with balance=F, logfc.threshold=0.14, max.genes=2000.

*Identification of cycling cells*

Cycling cells were identified as described in Neftel et al. (18). Enrichment scores were generated for each malignant cell in the same manner as in the previous section using two cell cycle signatures, one corresponding to G1/S and the other to G2/M. Cycling cells were identified by fitting each of the two cycling signature score distributions to a normal distribution and testing each cell’s significance using a threshold of 0.05. Cells that were significant for either of the two signature scores were labeled as cycling cells.

*Identification of putative glioma stem cells (GSCs)*

Malignant cells were labeled as GSCs based on nine definitions/gene signatures from seven publications (19–25) which we assigned abbreviated labels based on the specific phenotype and/or study: Bhaduri oRG, Xie qCSC, Couturier GPC, Wang mGSC, Antonica qPROM1, Bhaduri GSC, Suva TFs (Transcription Factors), Suva review, and Wang pGSC. Within Seurat, a new metadata variable was created for each of these, and cells were classified as GSCs or non-GSCs for each definition. For three GSC definitions, positive cells were identified by simple subsetting as in the original studies (Antonica qPROM1: non-cycling PROM1+ cells; Bhaduri GSC: TLR4-, SOX2+, and PROM1+ or FUT4+ or L1CAM+; Suva TFs: POU3F2+, SOX2+, and OLIG2+ cells, with SALL2 omitted given it was virtually not expressed in our dataset). For the remaining six GSC signatures, positive cells were identified using the method described in Xie et al. (20) briefly, the R package AUCell (26) was used to automatically identify positive and negative cell populations based on predicted activity of a specified gene signature, and this was performed for each of the six putative GSC gene signatures (lists available in Table S4).

*Cell-cell communication analysis*

Cell-cell communication analysis was performed using the CellChat package (27) with the functions and parameters recommended by the developers in their vignettes. This analysis was performed separately within each tumor. Malignant and non-malignant cell objects were merged, and cell type annotations were retained from the previous analyses except for tumor cells (analyzed as one “Glioma” group) and microglia/macrophages (analyzed as one “Myeloid” group). Each dataset was then split by region, and CellChat analysis was run using the CE and NE subsets separately, with raw.use=F (to ameliorate effects of dropout/sparsity) and population.size=F (to analyze communication independent of cell type composition). Differential analysis was then performed to compare CE and NE within each tumor. Only cell types present in both regions (at least 3 cells) were retained in the analysis. For cell type-level comparisons, both interaction count (Fig. S6C-E) and weight (Fig. 6) were analyzed. For pathway-level comparisons between NE and CE, the two quantitative metrics used were functional wiring (Fig. 7A, Fig. S7) and information flow (Fig. 7B-D, Fig. S8). To visualize overlapping findings across patients (Fig. 7, Fig. S6B), UpSet plots were generated using a custom script that utilizes a modified version of the R package ComplexHeatmap (28) that adds the ability to output overlapping pathway names next to the horizontal bar graphs. UpSet plots were chosen as an alternative to 5-group Venn diagrams for ease of visualization and interpretation.
